# Comparative analysis of piRNA sequences, targets and functions in nematodes

**DOI:** 10.1101/2022.06.26.497686

**Authors:** Hannah L. Hertz, Benjamin Pastore, Wen Tang

## Abstract

Piwi proteins and Piwi-interacting RNAs (piRNAs) are best known for their roles in suppressing transposons and promoting fertility. Yet piRNA biogenesis and its mechanisms of action differ widely between distantly related species. To better understand the evolution of piRNAs, we characterized the piRNA pathway in *C. briggsae*, a sibling species of the model organism *C. elegans*. Our analyses define 25,883 piRNA producing-loci in *C. briggsae*. piRNA sequences in *C. briggsae* are extremely divergent from their counterparts in *C. elegans*, yet both species adopt similar genomic organization that drive piRNA expression. By examining production of Piwi-mediated secondary small RNAs, we identified a set of protein-coding genes that are evolutionarily conserved piRNA targets. In contrast to *C. elegans*, small RNAs targeting ribosomal RNAs or histone transcripts are not hyper-accumulated in *C. briggsae* Piwi mutants. Instead, we found that transcripts with few introns are prone to small RNA overamplification. Together our work highlights evolutionary conservation and divergence of the nematode piRNA pathway and provides insights into its role in endogenous gene regulation.

## INTRODUCTION

Piwi proteins—members of the Argonaute family—and their piRNA cofactors are found in most of animals [1–3]. Studies from model organisms including *D. melanogaster*, *C. elegans* and mouse have provided key insights into piRNA biogenesis and functions. Genomic origins and sources of piRNAs are remarkably diverse across species. Single-stranded precursor transcripts are generated from piRNA-producing loci and then processed into mature piRNAs. Mature piRNAs are generally 21-35 nucleotides (nts) in length, preferentially start with a 5’ Uridine, and possess 2ʹ-O-methylated 3ʹ-ends [1–3]. The Piwi/piRNA pathway plays key regulatory roles in germline development and gametogenesis, hence is required for fertility [4–8]. So far, the best-established function of piRNAs across species is to maintain genome stability by silencing transposable elements in the animal germ line [9–11]. Yet accumulating evidence suggests that piRNAs regulate the expression of some endogenous transcripts and perform species-specific functions [12–17].

With powerful molecular and genetic tools, *C. elegans* becomes one of vital model systems in studying the piRNA pathway. A single functional *C. elegans* Piwi protein, named PRG-1, associate with piRNAs which are referred to as 21U-RNAs due to their strong propensity for a 5’- monoP uridine residue and length of 21-nts [18–22]. Two classes of piRNAs (type I and type II) were discovered [19]. Type I piRNAs are derived from two large clusters on chromosome IV [19–21]. The promoters of many type I piRNA genes contain an 8-nt consensus Ruby motif CTGTTTCA which is associated with chromatin factors including PRDE-1, SNPC-4, TOFU-4 and TOFU-5 [23–26]. Type II piRNA loci lack the Ruby motif and are associated with the promoters of many RNA polymerase II genes [19]. Both type I and II piRNAs are produced from 25- to 29-nt capped small RNA (csRNA) precursors [19]. To generate mature piRNAs, the 5’ cap and first two-nts of csRNAs are removed. Additional nucleotides are trimmed from their 3’ ends and 2’–*O*– methylation follows [27–30].

In *C. elegans*, most piRNAs lack obvious sequence complementarity to transposons [18–21]. By allowing mismatches, PRG-1/piRNAs recognize hundreds of germline transcripts and recruit RNA-dependent RNA Polymerases (RdRPs) to initiate the biogenesis of secondary siRNAs. These secondary siRNAs show the propensity for 22-nt length and 5’ guanine residues, thus are referred to as 22G-RNAs [31–34]. 22G-RNAs are loaded onto worm-specific Argonautes (WAGOs) that maintain and propagate epigenetic silencing [35–39]. Surprisingly, recent studies showed that *C. elegans* piRNA pathway promotes the expression of mRNAs and ribosomal RNAs by preventing unchecked amplification of 22G-RNAs [40–44]. It is unclear whether these observations are specific to *C. elegans* or can be applied to other nematode lineages or across phyla.

To better understand piRNA evolution and function, we characterized the piRNA pathway in *C. briggsae*, a nematode that diverged from *C. elegans* roughly 100 million years ago [45]. By sequencing small RNAs co-purified with *C. briggsae* Piwi protein, we identified 25,883 piRNA-producing loci and found that they are localized to at least three clusters on chromosome I and IV as well as promoter regions genome-wide [46]. We show that piRNAs induce the production of 22G-RNAs which are likely loaded onto WAGOs, but not onto CSR-1 Argonaute which are thought to protect endogenous germline genes from piRNA-mediated silencing [47, 48]. Furthermore, we identified 88 protein-coding genes that are targeted by both *C. elegans* and *C. briggsae* piRNA pathways. Although loss of Piwi leads to transgenerational infertility in *C. briggsae*, we did not detect hyper-accumulation of 22G-RNAs targeting ribosomal RNAs or histone mRNAs as observed in *C. elegans prg-1* mutants [41–44]. Upon loss of piRNAs, 22G-RNAs are overamplified preferentially from the transcripts possessing few introns. Altogether our comparative analyses revealed both conserved and species-specific features in the nematode piRNA pathways.

## RESULTS

### Identification and characterization of piRNA-producing loci in *C. briggsae*

The nematodes *C. briggsae (Cbr)* and *C. elegans (Cel)* diverged from a common ancestor tens of million years ago [45]. *C. briggsae* expresses a single Piwi protein: Cbr-PRG-1 (WormBase ID: WBGene00032975). Using CRISPR/CAS9 genome editing, we introduced 3xflag sequences to the genomic locus of *Cbr-prg-1*. Western Blotting analysis confirmed the expression of 3xFLAG::Cbr-PRG-1 protein (Supplementary Fig. 1A). To identify piRNAs in *C. briggsae*, we enriched 3xFLAG::Cbr-PRG-1 by immunoprecipitation (IP) and deep-sequenced the co-purified small RNAs, then aligned sequencing reads to the *C. briggsae* genome (WormBase release WS279). From two biological replicates, small RNAs from many distinct genomic loci were significantly enriched in Cbr-PRG-1 IP (IP/input ≥ 4-fold; p-value < 0.05, Two-tailed t-test) (Fig. 1A). Similar to their counterparts in *C. elegans*, the majority of Cbr-PRG-1 bound small RNAs are 21 nucleotides in length and exhibit a 5’ terminal uridine (Fig. 1B). We thus refer to them as 21U-RNAs. Our analyses defined in total 25,883 piRNA-producing loci (Supplementary Table S1), a number much larger than that previously estimated [46]. We found examples of discrete piRNA-producing loci (Supplementary Fig. 1B) as well as overlapping piRNA-producing loci (Supplementary Fig. 1C). To further validate authenticity of piRNA genes, we generated a presumptive null allele of *Cbr-prg-1* that bears a 5-nt deletion downstream of the start codon and shifts the open reading frame [15]. Consistent with previous findings that the stability of piRNAs and Piwi protein are co-dependent in *C. elegans*, the overall abundance of 21U-RNAs in the *Cbr-prg-1* mutant animals was reduced to ∼1% of wild-type (Fig. 1C).

**Figure 1:**
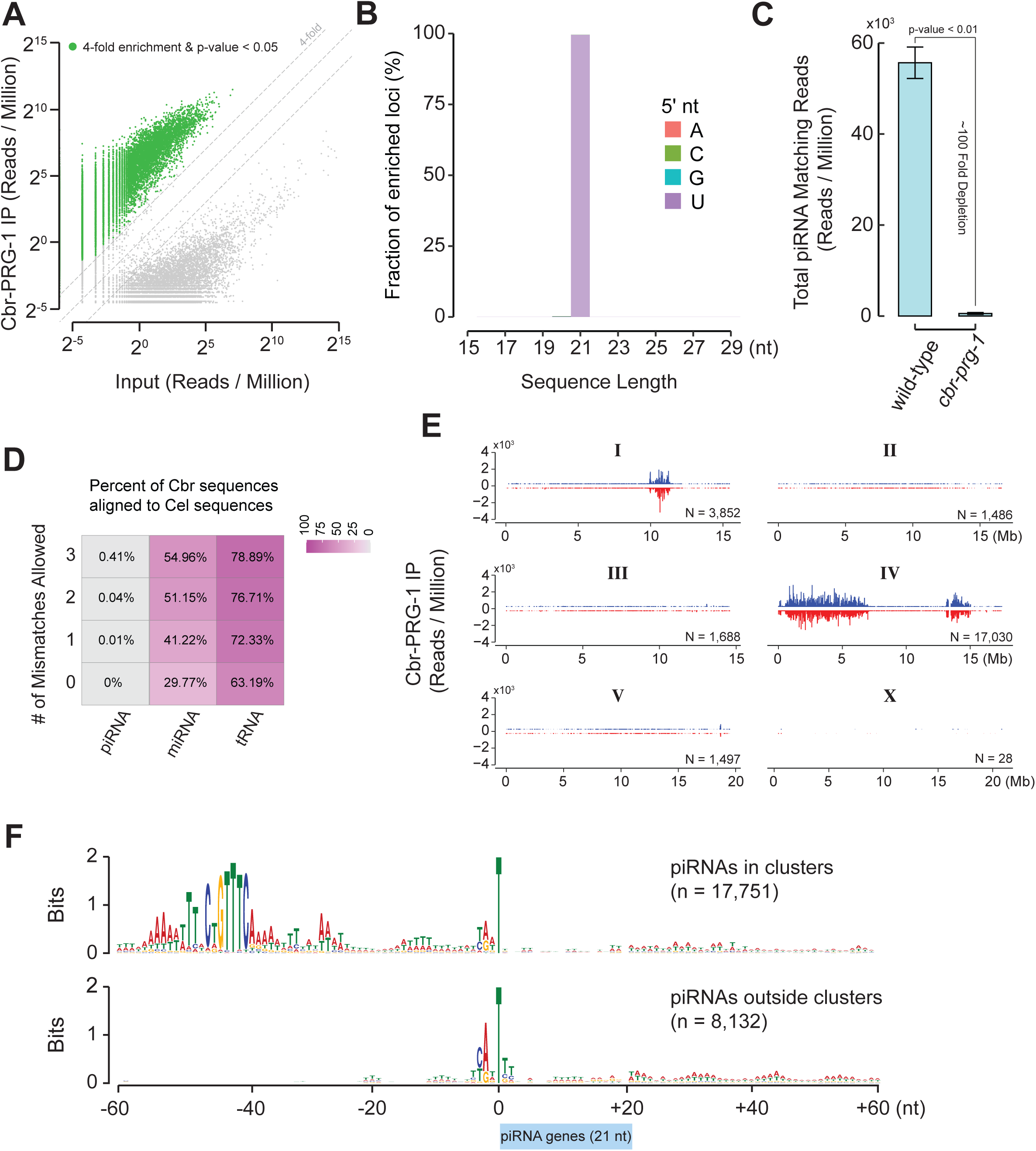
Identification and Characterization of piRNA-producing loci in *C. briggsae*. (A) Scatter plot showing the normalized abundance of sequence reads in 3XFLAG::Cbr-PRG-1 IP and Input samples. Data are displayed as an average of two biological replicates. Green dots mark reads 4-fold enriched in IP samples with a two-tailed t-test p-value < 0.05. Dotted grey diagonal lines represent 4-fold enrichment, no change in abundance, and 4-fold depletion, respectively. (B) Barplot showing the length and first nucleotide distribution of enriched reads from 3XFLAG::Cbr-PRG-1. Data are displayed as the fraction of enriched reads based on RPM from two averaged 3XFLAG::Cbr-PRG-1 IP biological replicates. (C) Barplot showing the overall abundance of C. *briggsae* piRNAs as determined by small RNA sequencing in wild-type and *prg-1* mutant animals. The error bars represent the mean ± the standard deviation of two biological replicates. A Two-tailed t-test was used to derive p-values. (D) Heatmap showing the percentages of C. *briggsae* piRNAs, miRNAs, and tRNAs mapping to a reference of *C.* elegans piRNAs, miRNAs, and tRNAs with 0, 1, 2, or 3 mismatches, respectively. (E) Barplots showing the location and abundance of piRNAs on each chromosome in C. *briggsae*. Blue bars represent piRNA loci residing on the forward strand of DNA, red bars represent loci residing on the reverse strand. The height of the bar reflects the abundance of each loci as an average of two PRG-1 IP biological replicates. 302 piRNA genes are mapped to contigs that do not have genomic coordinates and therefore are not presented in this panel. (F) The nucleotide preference within a 60-nucleotide window up- and down- stream of annotated piRNA genes that reside within and outside of piRNA clusters on chromosomes I and IV.

We assessed the sequence conservation of *Cbr*-piRNAs (n=25,883) and *Cel-*piRNAs (n=15,363) in comparison with the sequence conservation of miRNAs and tRNAs. Consistent with previous findings [46], 0%, 0.01%, 0.04%, and 0.41% of *C. briggsae* piRNAs had potential homologs in *C. elegans* when allowing zero, one, two, or three mismatches respectively (Fig. 1D). miRNAs are ∼22 nt small RNAs that regulate mRNA stability and suppress mRNA translation in diverse organisms [49]. *C. briggsae* miRNAs (n=131) and *C. elegans* miRNAs (n=368) exhibited stronger sequence conservation as compared to piRNAs. 29.77%, 41.22%, 51.15%, and 54.96% of *C. briggsae* miRNAs had potential homologs in *C. elegans* when allowing zero, one, two, or three mismatches respectively (Fig. 1D). The tRNA group displayed strongest sequence conservation among three non-coding RNA categories. 63.19%, 72.33%, 76.71%, and 78.89% of *C. briggsae* tRNAs had homologs in *C. elegans* when allowing zero, one, two, or three mismatches respectively (Fig. 1D). These findings indicate that in contrast to miRNAs and tRNAs, sequences of nematode piRNAs are poorly conserved. Since individual transcripts in the *C. elegans* germ line are often targeted and silenced by multiple piRNAs [33], it is possible that sequences of individual piRNAs are not subject to positive selection.

We next examined the distribution of piRNA-producing genes throughout *C. briggsae* genome. 17,751 piRNA-producing loci were concentrated in three large genomic clusters (9.8∼12.0 Megabase (Mb) on chromosome I, and 0.02∼7.1 Mb and 13.0∼15.0 Mb on chromosome IV) on both Watson and Crick strands (Fig. 1E and 1F) [46]. We also found that 8,132 piRNA-producing loci were widely distributed across chromosome I, II, III, IV and V, but scarcely present on chromosome X (Fig. 1E and 1F). piRNA within the clusters and outside the clusters bore a resemblance to type I and type II piRNAs discovered in *C. elegans* [19]. For example, similar to *C. elegans* type I piRNA genes, *C. briggsae* piRNA-producing loci within the clusters on both chromosome I and IV shared the upstream 8-nt Ruby motif CTGTTTCA (Fig. 1F) [20, 46, 50]. In contrast, Ruby motif was not found at the piRNA-producing loci outside of the clusters (Fig. 1F). Both types possessed a strong bias for pyrimidine (Y: cytosine/thymine) and purine (R: guanine/adenine) dinucleotide 3- and 2-nt upstream of T which corresponds to 5’ end of 21U-RNA. The Y/R dinucleotide is known as the initiator element for RNA Polymerase II transcription initiation [51, 52]. It is thus possible that the transcription of *C. briggsae* piRNA precursors initiate two nucleotides upstream of the 5’ end of mature piRNAs. Low melting temperature of the DNA-RNA hybrid induces RNA polymerase II pausing/termination at *C. elegans* piRNA genes [50]. Consistent with this notion, we found both classes of piRNAs A/T rich motifs downstream of 21U-RNA sequences which might serve as terminator signals (Fig. 1F). Finally, it’s worth noting that although the number of type I piRNAs is roughly twice as many as type II piRNAs, their overall abundance is ∼8-fold higher (Supplementary Fig. 1D). Altogether, our analyses revealed that *C. elegans* and *C. briggsae* piRNA sequences evolve rapidly despite conserved features of promoters and terminators.

### Synteny analysis of *C. elegans* and *C. briggsae* piRNA clusters

To investigate the evolution of piRNA clusters in nematode, we performed synteny analysis between *C. elegans* and *C. briggsae* genomes. Firstly, we determined syntenic regions and genomic collinearity on a genome-wide scale [53]. We further interrogated *C. briggsae* piRNA clusters (9.8∼12.0 Mb on chromosome I, and 0.02∼7.1 Mb and 13.0∼15.0 Mb on chromosome IV) and *C. elegans* piRNA clusters (4.75∼7.5 Mb and 13.5∼17.5 Mb on chromosome IV) (Fig. 1E) [18–22]. Two *C. elegans* piRNA clusters are syntenic with two regions within the single piRNA cluster at the left end of *C. briggsae* chromosome IV (Fig. 2A) [46]. Notably, these two regions are separated by a ∼1 Mb DNA segment. The segment is part of the piRNA cluster in *C. briggsae*, but its syntenic region is not part of the piRNA cluster in *C. elegans* (Fig. 2A). *C. briggsae* piRNA clusters on chromosome I (9.8∼12.0 Mb) and IV (13.0∼15.0 Mb) are in synteny with multiple regions at *C. elegans* chromosome I and IV, respectively (Fig. 2A). However, none of these syntenic regions robustly produce piRNAs in *C. elegans*. The synteny analysis thus revealed some degree of conservation in the genomic location of piRNA clusters between *C. elegans* and *C. briggsae*.

**Figure 2:**
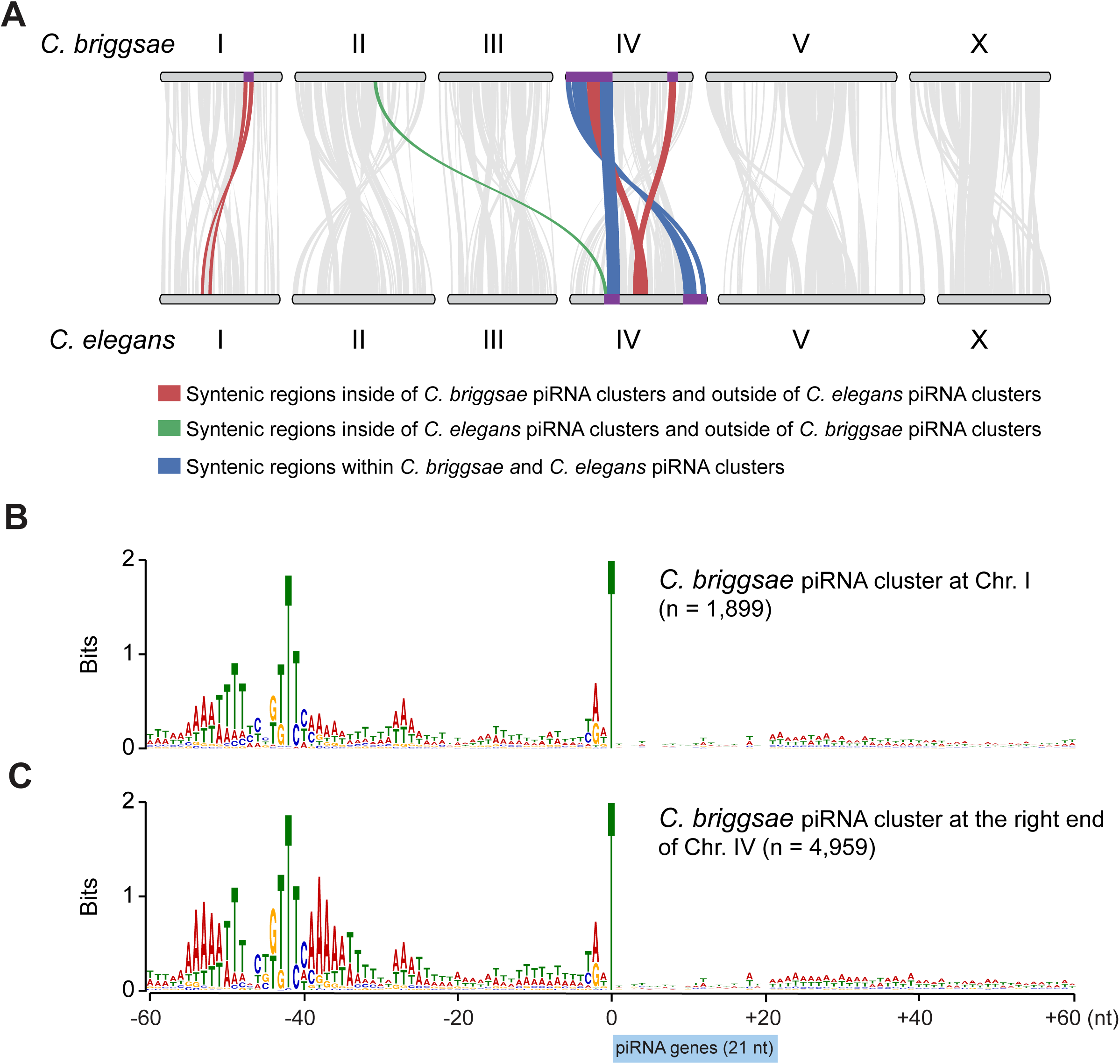
Evolution of *C. elegans* and *C. briggsae* piRNA clusters. (A) Synteny analysis between *C. briggsae* and *C. elegans*. Regions of syntenic collinear gene orthologs was found. Grey lines indicate regions of synteny between the two organisms. Red lines denote syntenic regions that reside within *C. briggsae* piRNA clusters. Green lines represent syntenic regions within *C. elegans* piRNA clusters. Blue lines indicate syntenic regions that reside within both the *C. briggsae* and *C. elegans* piRNA clusters. The location of *C. briggsae* and *C. elegans* piRNA clusters are highlighted in purple color on chromosomes I and IV. (B-C) The nucleotide composition within a 60-nucleotide window upstream and downstream of annotated piRNA genes that reside piRNA clusters on chromosome I (9.8∼12.0 Mb) and IV (13.0∼15.0 Mb), respectively.

Since *C. briggsae* genome possesses two species-specific piRNA clusters, we wish to examine whether piRNA-loci within these two clusters contain species-specific motifs. Similar to genome-wide analysis (Fig. 1F), both YR dinucleotide and Ruby motif were found at upstream of 21U-RNA sequences (Fig. 2B and 2C). Curiously, piRNA-producing genes, particularly ones at the *C. briggsae* piRNA cluster (chromosome IV 13.0∼15.0 Mb), exhibit strong A/T rich motifs upstream and downstream of the Ruby motif (Fig. 2C). The function of A/T rich motifs are unclear and require further investigation.

### Loss of Piwi causes progressive sterility in *C. briggsae*

Disruption of the piRNA pathway invariably causes sterility of animals examined to date [4, 7, 8, 54, 55]. In *C. elegans*, *prg-1* mutant animals do not become sterile immediately, but rather they become less fertile over generations, a phenotype associated with germ cell mortality [56]. We wondered if loss of *prg-1*/piRNAs in *C. briggsae* renders animals progressively sterile. To this end, we attempted to measure the brood size of wild-type and *Cbr-prg-1* mutant animals. Despite reported successes [57, 58], we failed to accurately measure the brood size of *C. briggsae* strains because of their burrowing behavior and avoidance to bacterial lawns. To tackle this issue, we decided to assess the fertility in the *Cbr-unc-119* mutant background. The *unc-119* gene encodes a lipid-binding chaperone expressed in the nervous system [59]. *unc-119* mutant animals exhibit slow movement [59], making it easier to measure the brood size. We outcrossed *Cbr-prg-1* with the *Cbr-unc-119* strains, freshly generated the *Cbr-unc-119* and *Cbr-prg-1*; *Cbr-unc-119* double siblings and conducted brood size measurement over generations. The fecundity of the *Cbr-unc-119* strain is maintained over generations. In contrast, *Cbr-prg-1*; *Cbr-unc-119* displayed progressive declines in fertility over generations (Fig. 3A).

**Figure 3:**
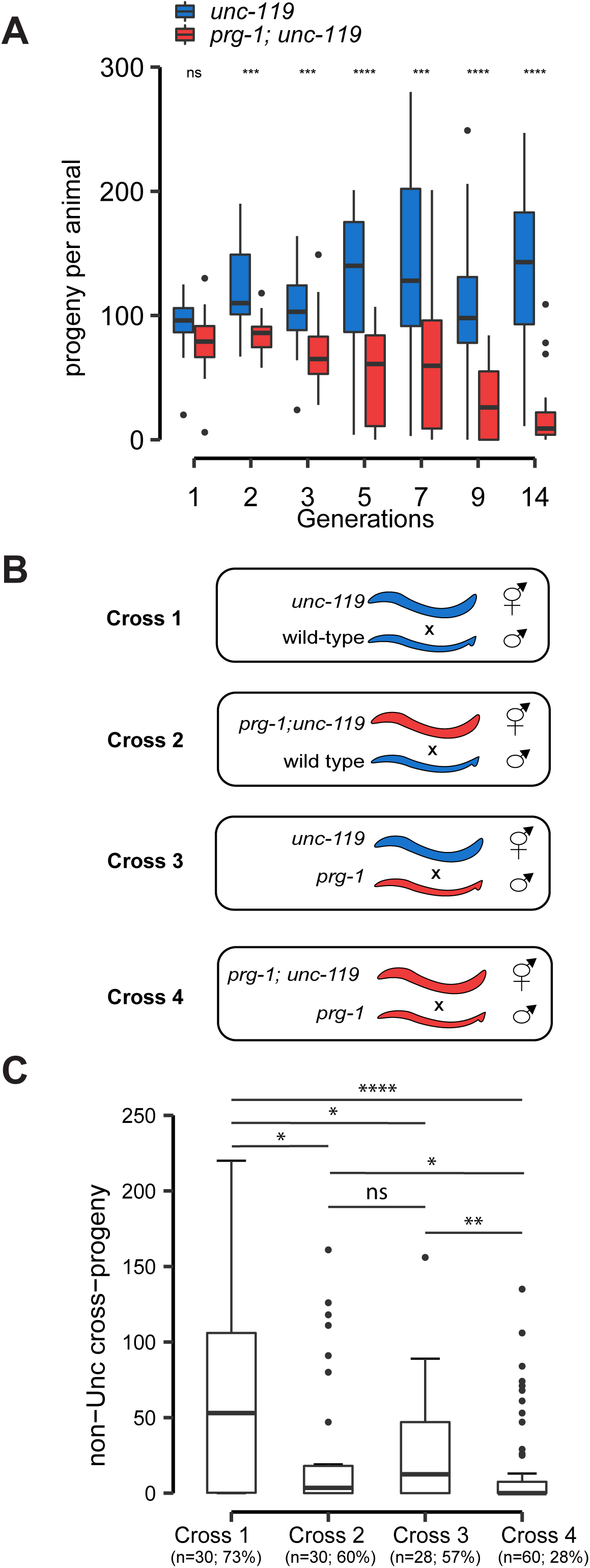
Loss of Piwi causes progressive sterility in *C. briggsae*. (A) Multigenerational brood size assay of *C. briggsae unc-119(nm67)* (blue and serves as wild-type control) and *prg-1; unc-119* (red) mutant animals at 25°C. The boxplots show the median (center line), 25 and 75^th^ percentile (box edges), and the whiskers indicate the median +/-1.5 x interquartile range. Fertility was assayed the following generations after outcross: 1 (wild-type n=15; *prg-1* n=15), 2 (wild-type n=17; *prg-1* n=19), 3 (wild-type n=22; *prg-1* n=25), 5 (wild-type n=22; *prg-1* n=25), 7 (wild-type n=23; *prg-1* n=24), 9 (wild-type n=21; *prg-1* n=21), and 14 (wild-type n=25; *prg-1* n=25). A Two-tailed Wilcoxon Rank Sum Test was used to compare fecundity of wild-type and *prg-1* mutants at each generation (ns = no significance, *p<0.05, **p<0.01, ***p<0.001, ****p < 0.0001). (B) Schematic of the cross strategy. *unc-119* mutant mothers were crossed with wild-type AF16 (cross 1) or *prg-1* mutant (cross 3) males. *prg-1*; *unc-119* mutant mothers were crossed to wild-type AF16 (cross 2) or *prg-1* mutant (cross 4) males. *prg-1* mutants are indicated by red. (C) Mating assay showing the median number of non-Uncoordinated (non-Unc) cross-progeny for each cross (cross 1, n=30; cross 2, n=30; cross 3, n=28; cross 4, n=60). The percentage of crosses with no non-Unc cross-progeny are shown. A Two-tailed Wilcoxon Rank Sum Test was used to compare total non-Unc cross-progeny, with the following significance indicators: (ns = no significance, *p<0.05, **p<0.01, ***p<0.001, ****p < 0.0001).

***C. briggsae*** hermaphrodites produce both sperm and oocytes. To determine whether the infertility of *Cbr-prg-1* mutants results from a defect in oogenesis and/or spermatogenesis, we measured male and female fertility based on four genetic crosses (Fig. 3B). When wild-type males mated with unc-119 hermaphrodites, the median brood size was 53 (as measured by counting the total number of non-Unc cross progeny) and 73% hermaphrodites produced non-Unc progeny (Fig. 3C). The brood size was decreased when wild-type males were crossed to *Cbr-prg-1* hermaphrodites. Reduction in the brood size was also observed when *Cbr-prg-1* males were crossed to wild-type hermaphrodites (Fig. 3C). When *Cbr-prg-1* males mated with *Cbr-prg-1* hermaphrodites, an additive fertility deficit was observed. Only 28% *Cbr-prg-1* hermaphrodites produced non-Unc progeny (Fig. 3C). These findings suggest that loss of *prg-1*/piRNAs causes impairment in oogenesis and spermatogenesis.

The Piwi/piRNA pathway is required for germ cell maintenance. For example, loss of Piwi is linked to increased germ cell apoptosis in *C. elegans*, zebrafish and mice [18, 21, 60]. To monitor apoptotic germ cell corpses, we imaged the germ line of late-generation (∼40^th^ generation) *Cbr-prg-1* mutants by staining with acridine orange (AO) [61]. AO is capable of entering into corpses upon increases in membrane permeability and staining DNA [61]. We found apoptotic corpses in gonads of both wild-type and *Cbr-prg-1* mutant strains, but did not observe any statistically significant differences between the two strains (Supplementary Fig. S2A and S2B), indicating loss of *prg-1*/piRNAs does not significantly enhance germ cell apoptosis in *C. briggsae* [18, 21, 60]. To monitor apoptotic germ cell corpses, we imaged the germ line of late-generation (∼40^th^ generation) *Cbr-prg-1* mutants by staining with acridine orange (AO) [61]. AO is capable of entering into corpses upon increases in membrane permeability and staining DNA [61]. We found apoptotic corpses in gonads of both wild-type and *Cbr-prg-1* mutant strains, but did not observe any statistically significant differences between the two strains (Supplementary Fig. S2A and S2B), indicating loss of *prg-1*/piRNAs does not significantly enhance germ cell apoptosis in *C. briggsae*.

### *C. briggsae* piRNAs induce production of 22G-RNAs targeting mRNAs

In *C. elegans*, the PRG-1/piRNAs complex recognizes germline transcripts and recruits RNA-dependent RNA Polymerases (RdRPs) for the production of 22G-RNAs which are loaded onto Worm-specific Argonautes (WAGOs) that maintain and propagate epigenetic silencing [32, 35–39]. *C. briggsae* genome contains at least 10 WAGO genes and 4 RdRP genes (WBGene00038666/*Cbr-rrf-1*, WBGene00024074/*Cbr-rrf-3*, WBGene00023729/*Cbr-ego-1*, and WBGene00026758) [46]. We wondered if *Cbr-PRG-1*/piRNAs induce the production of WAGO 22G-RNAs. Two computational analyses were performed to test this idea. In the first analysis, we predicted piRNA target sites in silico and measured 22G-RNA coverage around putative target sites. 21U-RNA target sites were predicted by mapping *Cbr-*piRNA sequences to the *Cbr-* transcriptome allowing 0 mismatches in the seed region (position 2-8), 1 G:U wobble base-pair in the seed region, 3 mismatches outside the seed region (position 9-21), and 2 G:U wobble base-pairs outside the seed region. This analysis identified 4528 potential piRNA target sites on 3592 gene (1.26 target sites/gene). A 22G-RNA peak directly over the predicted piRNA-binding sites was detected (Fig. 4A). In addition, there was a ∼1.75-fold enrichment of 22G-RNA levels in wild-type as compared to *Cbr-prg-1* mutants within ±50 nt of predicted piRNA sites. No enrichment was detected in the negative control in which reverse complementary sequences of 21U-RNA sequences were used to predict target sites (Fig. 4B). When considering the least abundant (bottom 25%) piRNA species, we found a weak yet noticeable enrichment of 22G-RNAs in wild-type compared to *Cbr-prg-1* animals (Fig. 4C). When considering the most abundant (top 25%) piRNA species, we noticed a strong enrichment (2.6-fold) of 22G-RNAs in wild-type relative to the *Cbr-prg-1* animals (Fig. 4D). The global analysis was further confirmed by inspecting specific 21U-RNA/target interactions. For example, we observed production of 22G-RNAs that overlapped with or were adjacent to predicted piRNA target sites at WBGene00041650 and WBGene00087786 loci (Fig. 4E and 4F).

**Figure 4:**
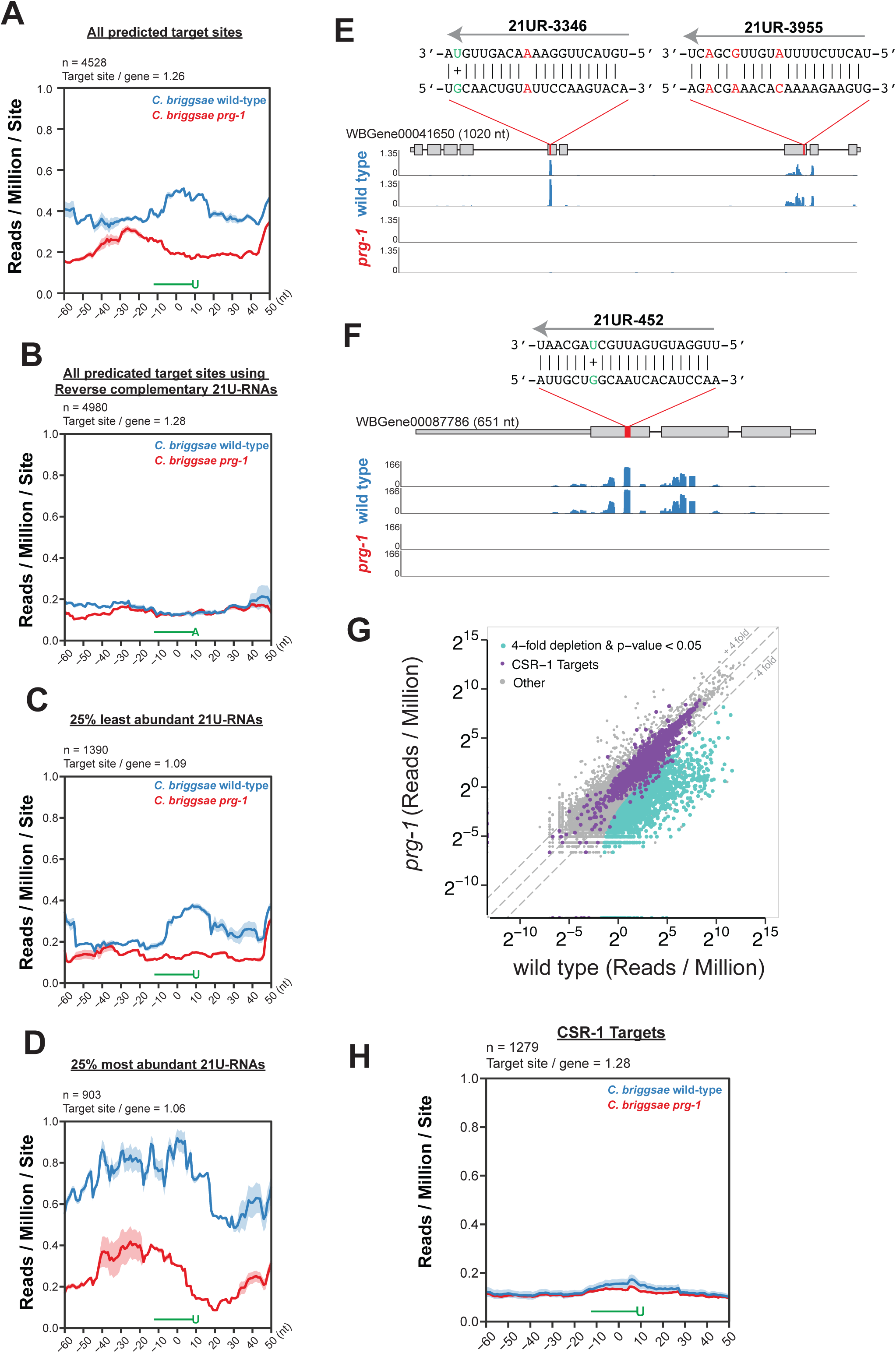
*C. briggsae* piRNAs induce production of 22G-RNAs mapped to endogenous mRNAs. (A-D). Line plots showing the coverage of 22G-RNAs within a 110 nt window centered on the 10^th^ nucleotide of targeting piRNAs for all predicted target sites (A), all predicted target sites using reverse complementary 21U-RNAs (B), target sites for the 25% least/most abundant piRNAs by IP RPM (C and D). Blue lines represent the coverage in wild-type and red lines indicate the coverage in *prg-1* mutants. Data are displayed as an average of two biological replicates, shaded regions indicate the standard deviation at each position plotted. Above each plot are the total number of piRNA target sites and the number of target sites per gene. (E-F) Browser view of 22G-RNA signal at predicted piRNA target sites on WBGene00041650 (E) and WBGene00087786 (F). Coverage of 22G RNAs along the transcript are shown for wild-type (blue) and *prg-1* mutants (red). A red bar within the gene model marks the precise 21UR-3346 and 21UR-3955 (E) and 21UR-452 (F) target sites. Above each red bar is a schematic illustrating the predicted base-pairing pattern between piRNAs and targets. Mismatched base-pairs are shown in red and G:U base-pairs are shown in green along with a + sign. (G) Scatter plot showing the abundance of 22G-RNAs targeting protein-coding genes, pseudogenes, lincRNAs, and transposons in *prg-1* and wild-type *C. briggsae*. Purple dots indicate CSR-1 targets annotated in [63]. Turquoise dots represent transcripts that are 4-fold depleted of 22G-RNAs (Two-tailed t-test, p-value < 0.05). (H) Line plot of the 22G-RNA levels within a 110 nt widow centered on the 10^th^ nucleotide of piRNAs on CSR-1 targets is shown. Data are displayed as an average of two biological replicates, shading above each line represent the standard deviation between the two biological replicates at each point. The number of target sites as well as the number of sites per gene are indicated above the plot.

As for a second analysis, we examined levels of 22G-RNAs targeting individual transcripts. In the *C. elegans* adult germ line, 22G-RNAs can be sorted into distinct pathways including the CSR-1 pathway and WAGO pathway [31, 62]. While *C. briggsae* small RNA pathways require further characterization, one study carefully characterized the Cbr-CSR-1 pathway and defined 4839 genes targeted by Cbr-CSR-1 [63]. When comparing 22G-RNA levels in *cbr-prg-1* mutants to wild-type animals, we found that the Cbr-CSR-1 targets remained largely unchanged (Fig. 4G). In contrast, many non-CSR-1 targets, which are presumably targeted by WAGOs, showed depletion of 22G-RNAs in *cbr-prg-1* mutants (Fig. 4G). In addition, we identified 1279 predicted 21U-RNA target sites on CSR-1 targets (Fig. 4H). Yet no significant enrichment of 22G-RNA levels was detected in wild-type as compared to *Cbr-prg-1*, consistent with the idea that CSR-1 and its small RNAs protect germline mRNAs from piRNA-mediated silencing (Fig. 4H) [47, 48]. In total, we defined 858 mRNAs, 3 lincRNAs, 45 pseudogenes, and 4 transposons as Cbr-PRG-1 targets whose 22G-RNAs are significantly depleted upon loss of *prg-1* (Cbr-prg-1/wild-type ≤ 4-fold, p-value < 0.05, reads per million (RPM) ≥ 5 in wild-type, Two-tailed t-test) (Supplementary Table S2).

### Conserved genes are targeted by piRNAs in *C. briggsae* and *C. elegans*

We next assessed the evolutionary conservation of PRG-1 targets between *C. briggsae* and *C. elegans*. ∼65% (13179/20157) of *C. briggsae* genes have orthologs in *C. elegans* and ∼63% (13179/20997) of *C. elegans* genes have orthologs in *C. briggsae* [63]. We cloned and deep sequenced small RNAs from *C. elegans* wild-type and *prg-1(tm872)* strains and identified 1776 Cel-PRG-1 targets based on the reduction of 22G-RNAs (Cel-prg-1/wild-type ≤ 4-fold, p-value < 0.05, RPM ≥ 5 in wild-type, Two-tailed t-test) (Supplementary Table S3). When comparing this set with Cbr-PRG-1 targets described above, we found that 58% (526/910) of Cbr-PRG-1 targets had orthologs, a percentage comparable to the genome wide average (Fig. 5A). A similar trend was observed in *C. elegans* in which 63% (1120/1776) of Cel-PRG-1 targets possessed orthologs (Fig. 5A). We next examined the genomic distribution of PRG-1 targets in *C. briggsae* and *C. elegans*. Cel-PRG-1 targets (n=1776) are relatively uniformly distributed on each chromosome, although they are slightly underrepresented on chromosomes V and X after normalized to number of genes per chromosome (Fig. 5B and Supplementary Fig. 3A). In contrast, Cbr-PRG-1 targets (n=910) are strongly enriched on X chromosome as compared to autosomes (Fig. 5C and Supplementary Fig. 3B), suggesting the *C. briggsae* piRNA pathway preferentially targets transcripts expressed from the sex chromosome.

**Figure 5:**
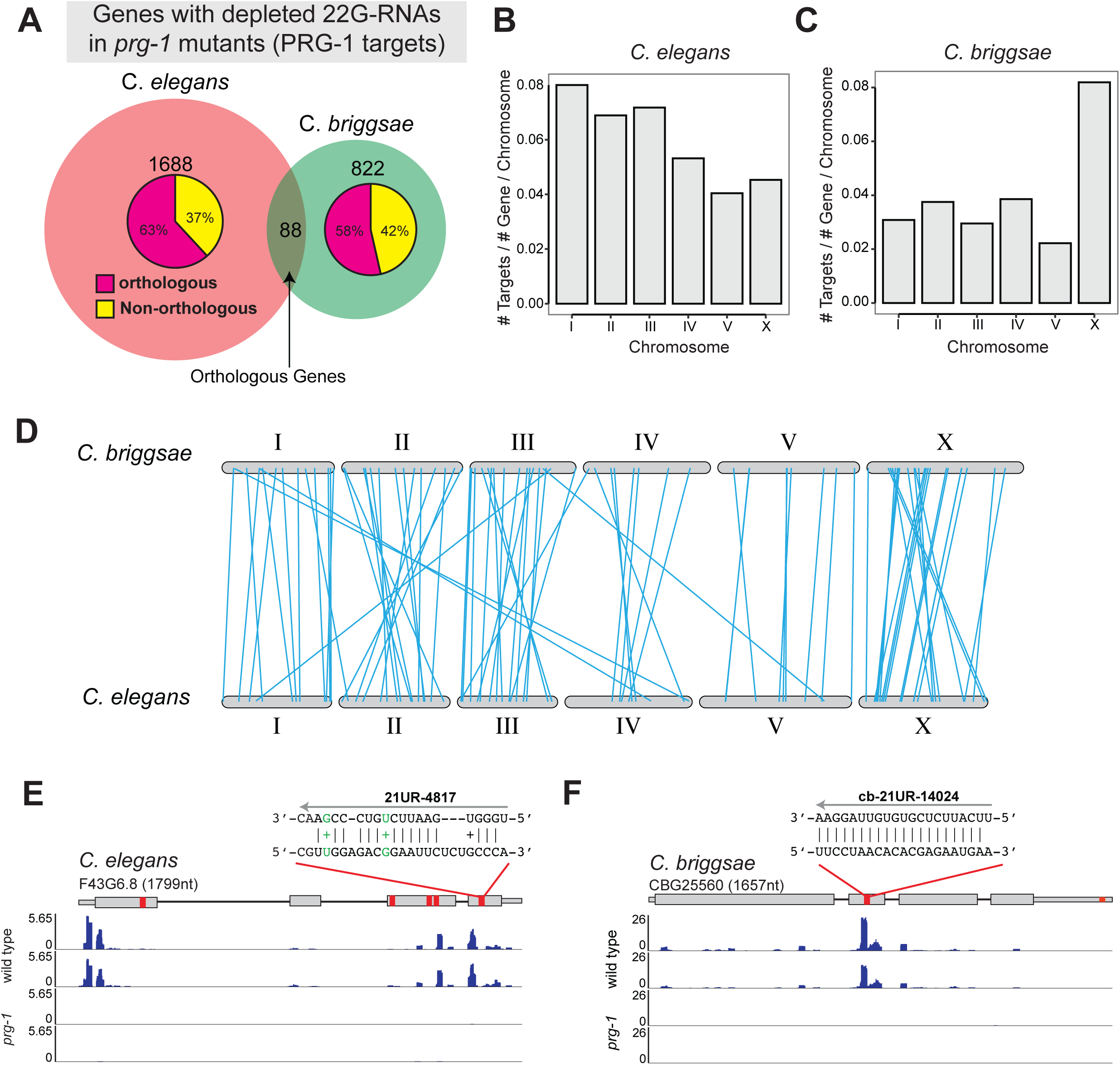
PRG-1 targets in *C. briggsae* and *C. elegans*. (A) Venn diagram showing the overlap of orthologous genes with decreased 22G-RNA abundance in C. *elegans* and C. *briggsae prg-1* mutants. The breakdown of non-overlapping genes with / without orthologs in the other species is shown as pie charts within each circle. (B-C) Barplots showing the number of PRG-1 targets per total number of genes per chromosome in *C. elegans* (B), and *C. briggsae* (C). (D) Karyotype map of *C. briggsae* and *C. elegans* showing the genomic location of conserved orthologous PRG-1 targets (blue lines). (E and F) Genome browser view of 22G RNA coverage near piRNA target sites for a conserved orthologous PRG-1 target pair: *C. elegans* transcript F43G6.8 (D), and the *C. briggsae* transcript CBG25560 (E). In each gene model red bars denote the location of putative piRNA target sites. *C. elegans* piRNA target sites were defined using *C. elegans* PRG-1 CLASH data [33]. *C. briggsae* piRNA target sites were predicted based on prediction (See Materials and Methods).

The best-established function of piRNAs is to recognize and silence transposable elements [9–11]. To our surprise, only 4 out of 208 annotated transposable elements (MARINER36, DNA2-19, HAT7, and LINE2B) are targeted by *C. briggsae* piRNAs (Supplementary Table S2). The *C. elegans* piRNA pathway targets 18 transposons (CELE42, CER13-I, CER6-I, Chapaev-2, HELITRON2, LINE2A, LINE2C, LINE2H, LONGPAL1, LONGPAL3, MARINER5, MSAT1, NDNAX2, NDNAX3, PALTTTAAA3, TC2, TURMOIL1, TURMOIL2) (Supplementary Table S3) [38]. This analysis revealed that distinct members of MARINER and LINE2 families are targeted by the piRNA pathways in both species.

Intriguingly, our analyses identified 88 evolutionarily conserved piRNA targets. These include *xol-1,* a master regulator of X chromosome dosage compensation and sex determination [15, 64] (Supplementary Table S4). Gene ontology (GO) analysis did not enrich any specific GO terms. We further examined the genomic location of 88 orthologous *C. elegans* and *C. briggsae* PRG-1 targets. They are distributed across autosomes and the X chromosome, but slightly depleted at chromosome V (Fig. 5D). 82/88 targets reside on the same chromosomes in both species, while 6/88 orthologs residing on different chromosomes of both species (Fig. 5D). Notably, even among conserved piRNA targets, PRG-1-induced 22G-RNA often target different regions of mRNAs. For example, *C. elegans* F43G6.8 (WormBase ID: WBGene00009660) encodes a putative metal ion binding protein. 22G-RNAs target its 5’ untranslated region (UTR) as well as the third and fourth exons of the transcript (Fig. 5E). In contrast, its *C. briggsae* ortholog, CBG25560 (WormBase ID: WBGene00086974), has abundant 22G-RNAs mapped to its second exons (Fig. 5E). 21U-RNAs and their target sites on *C. elegans* F43G6.8 were captured by a method known as crosslinking, ligation, and sequencing of hybrids [33, 65], while 21U-RNAs and their target sites of *C. briggsae* CBG25560 were predicted in silico (see above). In both cases, 22G-RNAs were found to be adjacent to putative piRNA target sites (Fig. 5E and 5F). Together, our analyses revealed a set of evolutionarily conserved PRG-1/piRNA targets which are targeted by different piRNAs at different regions.

### Loss of piRNAs leads to accumulation of 22G-RNAs at different sets of genes in *C. briggsae* and *C. elegans*

*C. elegans* piRNA pathway acts to prevent overproduction of 22G-RNAs [40–44]. In particular, upon loss of PRG-1/piRNAs, ribosomal RNAs (rRNAs), histone mRNAs, and additional transcripts are targeted by RdRPs which overamplify 22G-RNAs [40–44]. It has been postulated that mis-regulation of rRNA and/or histone leads to infertility in *C. elegans* [41, 42]. However, it is not clear why a specific set of RNAs are susceptible to 22G-RNA overproduction in *prg-1* mutants.

We reasoned that comparative analysis can offer insights into conserved features of misprocessed RNAs. Consistent with previous findings [40–43], we found 148 genes exhibited elevated 22G-RNA levels in *C. elegans prg-1* mutants (Cel-prg-1/wild-type ≥ 4-fold, RPM ≥ 5 in *prg-1*, p-value < 0.05, Two-tailed t-test). Among them, 69% (102/148) of these genes had orthologs in *C. briggsae* (Fig. 6A). Using the same parameters, 96 genes showed upregulated 22G-RNAs in *C. briggsae prg-1* mutants. Among them, 30% (29/96) of genes had orthologs in *C. elegans*, a much lower percentage than the genome-wide average (Fig. 6A). To our surprise, zero gene with 1:1 orthologs displayed elevated 22G-RNAs in *C. elegans* and *C. briggsae* (Fig. 6A). We examined 22G-RNAs that specifically target rRNAs and histone mRNAs. In *C. elegans*, levels of 22G-RNAs that are mapped to 39 histone loci were significantly elevated in *prg-1* mutants (Fig. 6B) [40, 41, 43]. And the median abundance of 22G-RNAs was upregulated 6.7-fold in *prg-1* mutants as compared to wild-type (Fig. 6B). In *C. briggsae*, however, most of histones did not display increased 22G-RNAs when *prg-1* is mutated (Fig. 6C). In *C. elegans*, 22G-RNAs from rrn-1.1, rrn-2.1 and rrn-3.1 were aberrantly accumulated in *prg-1* mutants (Fig. 6B) [42, 43]. Specifically, the median abundance of 22G-RNAs in *prg-1* was increased 5.5-fold relative to wild-type (Fig. 6B). In contrast, the level of 22G-RNA produced from rRNAs remained largely unchanged in *C. briggsae prg-1* mutant animals as compared to wild-type (Fig. 6C). Together, our analyses revealed that a unique set of RNAs accumulate 22G-RNAs in *C. briggsae* and *C. elegans* upon loss of *prg-1*.

**Figure 6:**
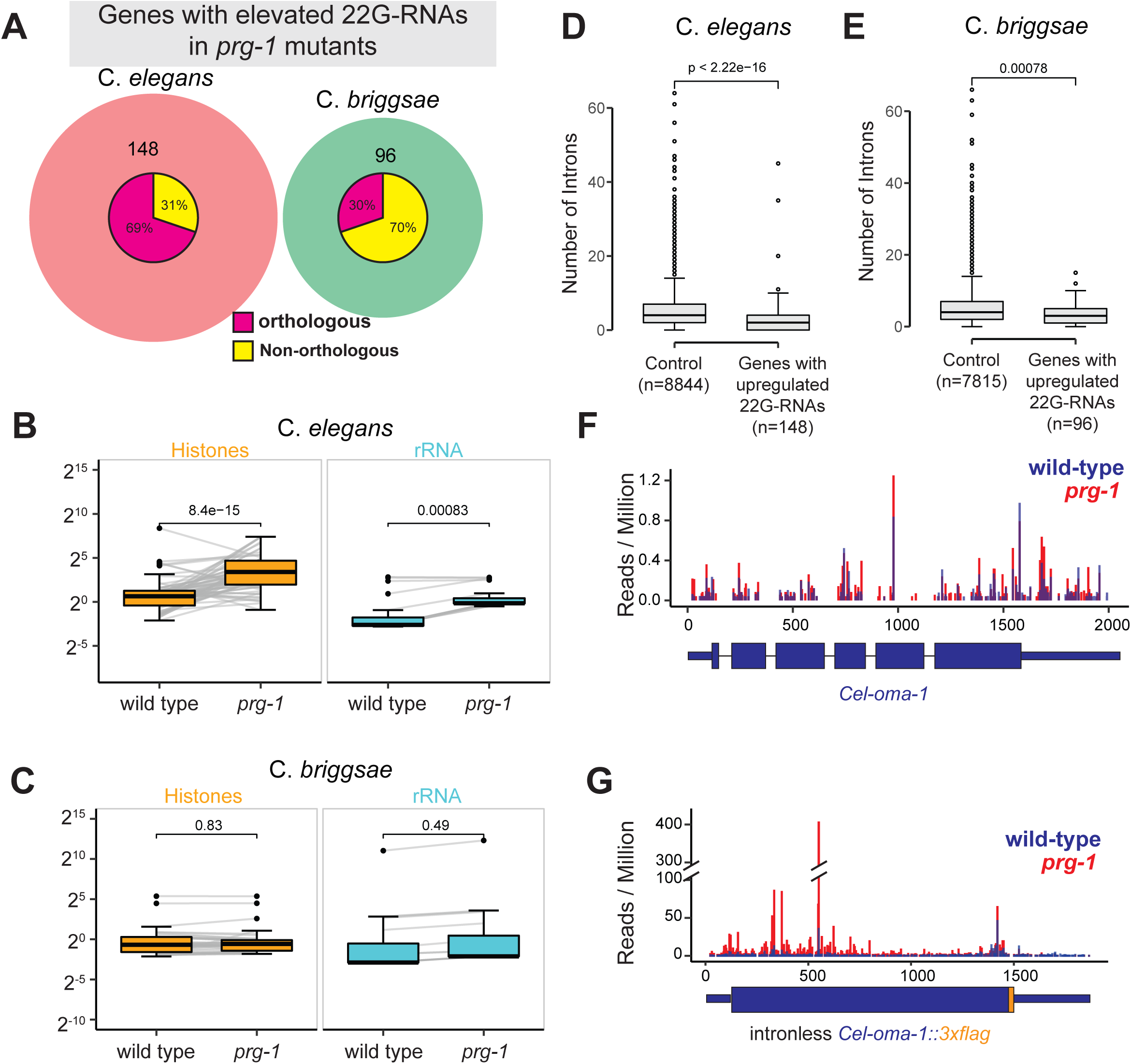
Characterization of genes that accumulate 22G RNAs in *prg-1* mutants. (A) Venn diagram showing the overlap of orthologous genes with increased 22G-RNA levels in C. *elegans* and C. *briggsae prg-1* mutants. The breakdown of non-overlapping genes with / without orthologs in the other species is shown as pie charts within each circle. (B-C) Boxplots showing the abundance of 22G-RNAs targeting histone genes and ribosomal RNAs in *C. elegans* (B) and *C. briggsae* (C). p-values were derived using a Two-tailed t-test. (D-E) Boxplots showing the number of introns in genes with accumulated 22G RNAs in *C. elegans* (D) and *C. briggsae* (E) *prg-1* mutants. The sample size of control and genes with upregulated 22G RNAs in *C. elegans* and *C. briggsae* are denoted in x-axis labels. Genes in the control set are expressed in the germ line and do not exhibit changes in 22G-RNA levels in *prg-1* mutants. p-values were derived using a Two-tailed Wilcoxon Rank Sum Test. (F-G) Histograms represent the position and abundance of 5’ end of 22G-RNAs targeting *C. elegans oma-1* transcript (F) or intronless *C. elegans oma-1::3xflag* (G) [71]. Blue bars represent reads from wild-type samples; red bars indicate reads from *prg-1* mutant samples.

### Relationship between splicing and hyper-accumulation of 22G-RNAs

We carried out comparative analysis to investigate common features of RNAs that are prone to 22G-RNA overamplification in the absence of piRNAs. 5’ UTR and 3’ UTR were examined due to their important regulatory functions [66, 67]. We compared transcripts exhibiting upregulated 22G-RNA levels to the control groups which are germline-expressed transcripts whose 22G-RNAs levels did not alter in *prg-1* mutants. No clear trend was detected regarding lengths of 5’ UTR and 3’ UTR in *C. briggsae* and *C. elegans* (Supplementary Fig. 4A-D). For example, 5’ UTR of RNAs (n = 148) with elevated 22G-RNAs are longer than the control group in *C. elegans*, while 5’ UTR of genes (n = 96) with elevated 22G-RNAs are shorter than the control set in *C. briggsae* (Supplementary Fig. 4A and 4C). The length of 3’ UTR in *C. elegans* and *C. briggsae* displayed an opposite trend (Supplementary Fig. 4B and 4D). Next, we examined introns, as there is evidence for the intimate interplay between pre-mRNA splicing and small RNA biogenesis [68–71]. We noticed that transcripts with upregulated 22G-RNA levels possess fewer introns in both *C. elegans* and *C. briggsae* relative to control groups which are germline-expressed transcripts whose 22G-RNAs levels did not upon loss of *prg-1* (Fig. 6D and 6E), indicating abundant introns or pre-mRNA splicing events may inhibit RdRP-mediated 22G-RNA production. Indeed, when further grouping *C. elegans* histone genes into intron-containing histones and intronless histones, we found that intronless histone group (n=64), but not the intron-containing histones (n=8), exhibited increased 22G-RNAs in *prg-1* mutants relative to wild-type (Supplementary Fig. 4E). In total, 39 histone mRNAs showed upregulated 22G-RNAs in *C. elegans prg-1* mutants (*prg-1*/wild-type ≥ 4-fold, reads per million (RPM) ≥ 5 in *prg-1*, p-value < 0.05, Two-tailed t-test) (Supplementary Table S3). Using the same parameters, however, we found that 0 histone genes exhibit elevated 22G-RNAs upon loss of *C. briggsae prg-1* (Supplementary Table S2).

To directly test the causal relationship between splicing and 22G-RNA synthesis, we assessed 22G-RNAs produced from the endogenous *C. elegans oma-1* gene and an intronless *oma-1* allele in which all intronic sequences were removed [71]. In strains expressing wild-type (intron-containing) *oma-1*, small RNA sequencing revealed comparable levels of 22G-RNAs between *wild-type* and *prg-1* mutants (Fig. 6F). 22G-RNAs were primarily mapped to exons, suggesting RdRPs use spliced *oma-1* mRNA as template to synthesize 22G-RNAs (Fig. 6F). In wild-type and *prg-1* mutant strains expressing intronless *oma-1*, abundant 22G were detected. In particular, loss of *prg-1* led to at least 5-fold increase in 22G-RNA level when compared to wild-type (Fig. 6G) [71]. Altogether our findings suggest the presence of functional introns—likely splicing itself or deposition of exon junction complex—prevents 22G-RNA overproduction in *prg-1* mutants (see Discussion below).

## DISCUSSION

In this study, we defined piRNA-producing loci and characterized piRNA functions in *C. briggsae*, a sibling species that separated from *C. elegans* about 100 million years ago [45]. Our comparative studies uncovered both conserved and species-specific features in the nematode piRNA pathways. Specifically, we identified over 25,000 C. *briggsae* piRNAs, a number that is comparable to that of their counterparts in *C. elegans* [18–21]. Although the architecture of piRNA genes and chromatin factors associated with piRNA clusters appeared to be conserved, piRNA/21U-RNA sequences have diverged rapidly (Figs. 1 and 2). Thus, piRNA sequences themselves are under little selective pressure. In contrast, the biogenesis mechanism and the number of piRNAs are under strong selection. Our findings raise some outstanding questions that require further investigation: 1) How do the rapidly evolving piRNAs fulfill their role in promoting fertility? 2) How are over 20,000 piRNAs-producing genes maintained over 100 million years of evolution? and 3) How do new piRNA genes arise? In flies, new piRNA species are produced when invading transposons are inserted into piRNA clusters and dispersed genomic regions [72, 73]. In worms, however, transposons are not enriched at piRNA clusters, nor did the majority of piRNAs display sequence complementarity to transposons. Thus, a yet undiscovered mechanism may be responsible for the generation of new piRNA-producing genes.

One interesting function of *C. elegans* piRNAs appears to be the recognition and silencing of foreign sequences [35–38]. By allowing mismatches between piRNAs and their target RNAs, piRNAs can recognize targets and induce 22G-RNA production by recruiting RdRPs (Fig. 4) [35–39]. Perhaps counterintuitively, the *C. elegans* piRNA pathway also prevents overamplification of 22G-RNAs [40–44]. Similarly, loss of *prg-1* in *C. briggsae* induces overproduction of 22G-RNAs targeting a set of mRNAs, although it is not clear whether excessive 22G-RNAs associate with Cbr-WAGOs or Cbr-CSR-1 (Fig. 6). In both organisms, RNAs with fewer introns are more susceptible for 22G-RNA overproduction. Removal of introns from *oma-1*, an endogenous gene, led to increased 22G-RNA levels when *prg-1* is mutated (Fig. 6G). These findings suggest that introns and/or efficient pre-mRNA splicing limits 22G-RNA overamplification. A connection between splicing and small RNA synthesis has been first demonstrated in yeast *Cryptococcus neoformans* and later found in nematode *C. elegans*, where stalled or inefficient splicing direct transcripts into the RNA interference (RNAi) pathway [69, 71, 74]. It is thought that some RNAi components surveil splicing to detect invasive sequences such as transposons and viral genomes. We envision that loss of *prg-1* frees up some RdRPs and WAGOs. It is possible that RdRPs intrinsically favor intronless and/or poorly spliced transcripts as templates to synthesize 22G-RNAs. Presumably, RNAs with fewer introns have fewer exon-junction complexes (EJCs). EJC are deposited onto RNAs during splicing, and plays key roles in their export, localization, and turnover [75]. A highly speculative hypothesis predicts EJC physically blocks the 3’ to 5’ polymerase activity of RdRPs. Alternatively, EJC may prevent RNAs from entering into mutator foci where 22G-RNAs are synthesized [76]. It is not clear why intronless histone mRNAs are immune to 22G-RNA overamplification in *prg-1 C. briggsae*. Previous studies reported a positive feedback loop in which 22G-RNAs initiate cleavage followed by pUGylation of RNAs and RdRPs use pUGylated RNAs as templates to synthesize more 22G-RNAs [77]. We noticed some basal levels of 22G-RNA in wild-type *C. elegans*, while 22G-RNAs targeting histone mRNAs are absent in wild-type *C. briggsae*. Perhaps *C. elegans* histone mRNAs, but not *C. briggsae* counterparts are primed for 22G-RNA overamplification. The presence of 22G-RNAs and/or pUGylated RNAs may be required for triggering 22G-RNA overproduction upon loss of *prg-1*/piRNAs.

Loss of *prg-1* in both *C. briggsae* and *C. elegans* leads to progressive sterility (Figure 3) [56]. Presumably, such sterility defect results from the loss or gain of 22G-RNAs for one or more transcripts. In *C. elegans*, the piRNA pathway prevents the overproduction of 22G-RNAs targeting rRNAs, histone mRNAs and additional transcripts [40–44]. It has been postulated that mis-regulation of rRNA or histone mRNAs contributes to *Cel*-*prg-1* infertility [41, 42]. However, loss of *Cbr-prg-1* does not lead to hyper-accumulation of 22G-RNAs targeting rRNAs, histone mRNAs or any *C. elegans* orthologs, suggesting that the 22G-RNA overamplification is not the evolutionarily-conserved mechanism for the infertility defect of Piwi mutants. Our analysis further revealed that *C. briggsae* piRNA pathway induces WAGO 22G-RNAs from hundreds of germline transcripts (Fig. 4). While CSR-1/22G-RNAs targets are highly conserved in *C. elegans* and *C. briggsae* [63], PRG-1/piRNA targets have diverged rapidly (Fig. 5). Nevertheless, 88 orthologous genes and members of MARINER and LINE2 transposon families were found to be targeted by both *C. briggsae* and *C. elegans* piRNA pathways (Supplementary Table S4). We envision that regulation of these evolutionarily conserved piRNA targets may promote fertility or contribute to other important physiological processes. Indeed, *xol-1*, one of the piRNA targets, is linked to worm dosage compensation and sexual development [15, 64]. Additional genetic and biochemical experiments will be required to determine physiological roles of other conserved piRNA targets.

Finally, significant progress has been made in understanding piRNA biogenesis and function using model systems including *D. melanogaster*, *C. elegans* and mice[1–3]. However, studies should not be limited to model organisms. In the past, next-generation sequencing technology has been employed to define piRNAs in non-model organisms [78–80]. With the advent of CRISPR genome editing tools [81], one can start interrogating piRNA targets and functions in non-model organisms. [78–80]. With the advent of CRISPR genome editing tools [81], one can start interrogating piRNA targets and functions in non-model organisms. Comparative genomic and functional studies will be essential to understand the rapidly evolving piRNA pathway.

## MATERIALS AND METHODS

### Worm strains

AF16 and N2 strain are reference strains for *C. briggsae* and *C. elegans* respectively. Worms were cultured according to standard methods at 20 °C unless otherwise indicated [82]. Mutant animals were generated using CRISPR editing or obtained from the CGC. All strains used in this study are listed in Supplementary Table S5.

### CRISPR genome editing

CRISPR/Cas9-generated strains were made as previously described [83]. In brief, single-stranded donor (2.2 µg) was used to introduce 3xFLAG to *prg-1*. Repair template donors were added to pre-assembled Cas9 ribonucleoprotein complex (5 µg Cas9, 2 µg gRNA, 1 µg tracrRNA) (IDT). Plasmid pRF4::rol-6(su1006) was used as a co-injection marker [83].

### Total RNA isolation

*C. briggsae prg-1* mutant and its wild-type siblings were grown continuously. Approximately 100,000 synchronized L1 animals at ∼10^th^ generation were obtained and plated on 135 mm NGM plates seeded with OP50 and grown at 20 °C until the animals reached gravid adult stage. Gravid adult populations were harvested using M9, and subsequently suspended in Trizol. Worms were lysed using the Bead Mill 24 homogenizer (Thermo Fisher Scientific). Bromochloropropane was added to the lysis to perform RNA extraction. Isopropanol was then used to precipitate RNA from the aqueous phase. Small RNAs were isolated from total RNAs using the miRVana miRNA isolation kit (Thermo Fisher Scientific) according to the manufacturers protocol.

### Small RNA sequencing

Small RNA samples from wild-type and mutants were first incubated with a recombinant 5’ polyphosphatase PIR-1 which removes the γ and β phosphates from 5′-triphosphorylated RNAs [84, 85]. The resulting monophosphorylated RNAs were ligated to the 3’ adaptor (5’rAppAGATCGGAAGAGCACACGTCTGAACTCCAGTCA/3ddC/3’, IDT) using T4 RNA ligase 2 in the presence of 25% PEG8000 (NEB) at 15 °C overnight. The 5’ adaptor (rArCrArCrUrCrUrUrUrCrCrCrUrArCrArCrGrArCrGrCrUrCrUrUrCrCrGrArUrCrU, IDT) was then ligated to the product using T4 RNA ligase 1 (NEB) at 15 °C for 4 hours. The ligated products were converted to cDNA using SuperScript IV Reverse Transcriptase (Thermo Fisher Scientific). The cDNAs were amplified by PCR using Q5 High-Fidelity DNA polymerase (NEB), and the libraries were sequenced on an Illumina Novaseq platform (SP 2 X 50 bp) at the OSU Comprehensive Cancer Center genomics core.

### Synteny analysis of piRNA clusters

MCscan and associated python scripts were used to compute and visualize regions of synteny between *C. briggsae* and *C. elegans* [53]. The MCscan pipeline uses bed annotations of protein-coding genes, and coding sequence (CDS) to find regions of synteny. The CDS fasta files were download from WormBase (WS279) for both *C. elegans* and *C. briggsae*. Gff3 files were converted to bed files for protein coding genes using custom python scripts. The synteny of piRNA clusters between *C. elegans* and *C. briggsae* was inferred from protein coding genes that reside within the piRNA clusters.

### Germline apoptosis and scoring

To quantify germline apoptosis, synchronized *C. briggsae* wild-type and *prg-1* mutant worms plated on Day 1 were grown at 20°C until Day 4. On Day 4, 50 worms per experiment were transferred to a fresh plate. 200 µL Acridine Orange (AO) stain (75 µg/mL in M9) was distributed evenly to the fresh plates and allowed to dry in the dark. After feeding on the AO-soaked bacterial lawn in the dark at 20°C for 1 hour, the worms were transferred to a clean plate seeded with *E. coli* and incubated in the dark for 2 hours to clear excess dye from the intestines. The worms were imaged under the 40x objective with fluorescence microscopy to quantify the number of apoptotic corpses per gonad arm. One gonad arm was scored per animal, and 33-40 animals scored per experiment.

### Multigenerational Brood Size Assays

*C. briggsae* wild-type (AF16) and *prg-1* mutant brood sizes were assayed at 25°C in an *unc-119(nm67)* background. After outcross, 25 independent lines for each strain were continuously grown each generation by transferring 4 L4 larvae per line to new plates. Brood Size assays were performed approximately every two generations. To assay brood size, 1 L4 larvae per line was placed singly on plates. The animals were transferred halfway through egg-laying and the total brood size for each animal was calculated by adding the progeny of the original and transferred plates.

### Mating assays

To test mother-dependent and/or father-dependent fertility defects of *C. briggsae prg-1* mutants, male stocks of *C. briggsae* wild-type AF16 or *prg-1* mutants were generated via self-cross and propagated for at least 12 generations. For each cross, 5 *unc-119(nm67)* hermaphrodites were plated with 10 males, with a total of 29-60 hermaphrodites per experiment. Animals were allowed to cross overnight at 20°C and single picked to new plates the next day. The mothers were transferred mid-egg laying and the total wild-type looking (non-Uncoordinated) cross progeny for each animal was calculated by adding the wild-type cross progeny from the original and transferred plates. Animals with no wild type looking progeny, either from unsuccessful crosses or fertility defects, were recorded as zero.

### Small RNA sequencing data analysis

Raw small RNA sequencing reads were parsed from adapters and low-quality reads using TrimGalore and assessed using FastQC. 15-40 nucleotide long reads were collapsed using custom shell scripts and aligned to the C. *elegans* or C. *briggsae* (WormBase release WS279) genome reference using Bowtie with the parameters -v 0 -m 1 -a --best --strata to obtain perfectly and uniquely mapped smRNA sequencing alignments [86]. Due to the repetitive nature of histone and rRNA genes, we realigned our smRNA sequencing reads to the genome using the Bowtie parameters -v 0 -m 1000 -a --best --strata to capture multimapping reads associated with these features [86]. Reads failing to align to the genome reference were realigned to a reference containing exon-exon junctions using the same Bowtie parameters. Exon-exon junction mapping reads were converted to genomic coordinates using custom python scripts and added to the genomic alignment. Sam alignment files were then converted to bed files using BEDOPS [87]. Following alignment, reads were normalized to the number of times mapped, and assigned to genomic features using BEDtools and custom python scripts [88]. Briefly, reads were filtered based on first nucleotide, length, strand orientation, 5’ to 5’ distance between read and feature using the following parameters: 1) *piRNAs.* Reads must map sense to piRNA loci, contain a 5’ U, 20-33 nt long and align to position 0,1 or 2 of annotated 5’ piRNA end; 2) *22G-RNAs.* Reads must map antisense to protein coding gene exon, pseudogene exon, lincRNA, rRNA, or annotated transposon. Only reads 21-23 nt in length starting with a 5’ G were considered.

Reads assigned to multiple genomic features were further processed using a hierarchical based filtering approach. Briefly, reads mapping to highly contaminating RNA species such as ncRNA, tRNA, snRNA, snoRNA, and rRNA were given priority over other features. Read counts for other multi-feature mapping reads were split equally amongst the number of features mapped. In addition to genome mapping reads were mapped to RepBase transposon consensus sequences and filtered using the same parameters described above [89]. Of note, rRNAs are poorly annotated in the C. *briggsae* reference genome, thus, smRNA sequencing reads from C. *briggsae* wild-type and *prg-1* mutants were aligned to the C. *elegans* reference to quantify 22G-RNAs targeting rRNAs.

### Defining *C. briggsae* piRNA-producing genes

Reads from IP and input samples were processed using TrimGalore and assessed using FastQC. Processed reads 15-40 nt in length were collapsed using custom shell scripts and aligned to the C. *briggsae* genome reference (WS279) using the Bowtie with the parameters -v 0 -m 1 [86]. Sam alignments were converted to bed files using BEDOPS [87]. Following alignment, IP reads with a count of less than 2 were removed. Aligned reads passing filters were then grouped based on their strand and 5’ nucleotide position. For a given genomic position/strand group reads were collapsed into unique loci; the length of the locus was determined as the length of the most abundant read within each group. Collapsed loci were then compared between IP and input samples by calculating fold change using the 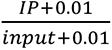, for both replicates of IP, as well as derivation of p-values using a Two-tailed t-test. Enriched loci were defined by 4-fold enrichment in both IP samples, and a p-value of less than 0.05. The 5’ nucleotide and length of enriched loci were plotted to elucidate the length and 5’ nt preference of C. *briggsae* piRNAs. Following this analysis, loci that are 21-nt in length starting with 5’ U were considered as C. *briggsae* piRNA.

### Motif Analysis

Genomic sequences 60 nts up and down stream of piRNA 5’ ends were isolated using BEDtools [88]. MEME motif analysis software was then used to construct sequence logos [90].

### 21U-RNA target site prediction

21U-RNA target sites were predicted using an analysis approach similar to that described [91]. Briefly, 21U-RNA sequences were aligned to a transcriptome reference containing protein coding genes, pseudogenes, lincRNAs and RepBase transposon sequences allowing up to 6 mismatches with BWA [89, 92]. Aligned sequences were then further processed to annotate the number of seed mismatches, seed G:U base-pairs, non-seed mismatches, non-seed G:U base-pairs, of each piRNA-target interaction using custom python scripts. For downstream metagene analysis piRNA target sites with less than or equal to 1 seed G:U base-pair, 3 non-seed mismatches, and 2 non-seed G:U base-pairs were considered.

### Density of 22G-RNAs

Processed collapsed smRNA reads were aligned to the same transcriptome reference used in 21U-RNA target site prediction using Bowtie. Sam files were converted to bedfiles using BEDOPS [87]. Antisense mapping 21-23 nt reads starting with G were then filtered from bed files. The per-base 22G-RNA coverage, normalized to total genome matching reads, was then calculated for all transcripts using custom python scripts. Regions of transcripts within 120 nucleotides of a piRNA target site were isolated and aggregated based on the relative position to the 10^th^ nucleotide of the piRNA. Plots showing the coverage of 22G-RNAs are of two averaged biological replicates and were drawn in R. piRNA target sites were assigned to genomic coordinates by mapping the target sequence to the genome using Bowtie.

### Definition of intronless and intron-containing histone genes

Histone genes were grouped into two categories based on the presence or absence of introns according to WormBase (WS279) annotation. Although *his-57* is annotated to have three isoforms (one is intronless and the other two contains introns), 22G-RNAs are found to be produced exclusively from the intronless isoform. Thus we reasoned that his-57 is likely mis-annotated and considered it as an intronless histone gene.

### Quantification and Statistical Analyses

All statistical tests used in this study are listed in the figure legends. Statistical tests and plots were preformed using custom R scripts. The major R packages used in the analysis of the data were dplyr, ggplot2, and other packages from Tidyverse [93].

### Data availability statement

The *C. briggsae* PRG-1 IP, as well as small RNA sequencing data from wild type and *prg-1* mutant *C. elegans* and *C. briggsae* strains are available at GEO under the accession number GSE203297. CSR-1 IP data are available at NCBI SRA under the accession number SRP021463 [63]. CLASH sequencing data used in this study to define piRNA target sites are available at NCBI SRA under the accession number SRP131397 [33]. All tables and metadata files are deposited to and are available at Dryad (https://datadryad.org/stash/share/JG5_VTsoNyi_62H03WHjnHH1wD2ZwZMLWpL4SnD1JTA)

## ACKNOWLEDGMENTS

We thank D. Schoenberg for discussion and comments; S. Tu for assistance in initial data analysis; the Neuroscience Imaging Core for instruments (S10OD026842); the OSU Comprehensive Cancer Center genomics core for Illumina sequencing. Some of the *C. elegans* and *C. briggsae* strains were provided by the Caenorhabditis Genetics Center supported by National Institutes of Health (P40 OD010440).

## Supplementary material

**Supplementary Figure 1.**
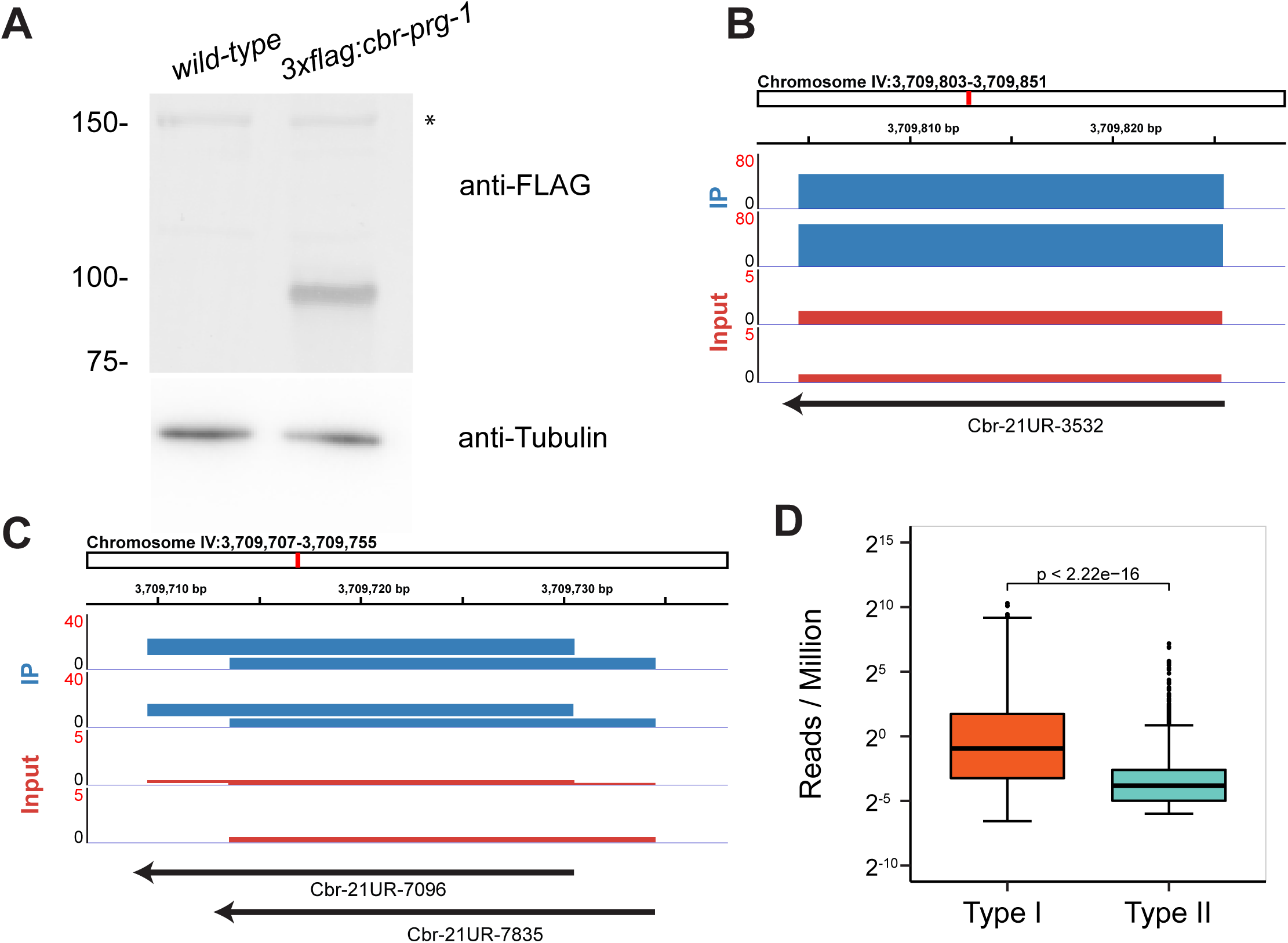
Expression of 3xFLAG::PRG-1 and piRNAs. (A) Western blotting analysis of 3xFLAG::PRG-1 in wild-type *C. briggsae* and *C. briggsae* expressing 3 x FLAG::PRG-1 with anti-FLAG antibody (top) and anti-Tubulin antibody (bottom). The asterisk denotes a background band. (B-C) Genome browser view of discrete and overlapping C. *briggsae* piRNA-producing loci. The scale of Cbr-PRG-1 IP (blue) and input (red) was adjusted so that reads from individual samples could be visualized. (D) Boxplot displaying the abundance of Type I (within clusters) and Type II (outside clusters) piRNA in C. *briggsae* as determined by small RNA sequencing. The thick bar within each box represents the median normalized piRNA abundance in each category. A Two-tailed Wilcoxon Rank-Sum test was used to derive p-values.

**Supplementary Figure 2:**
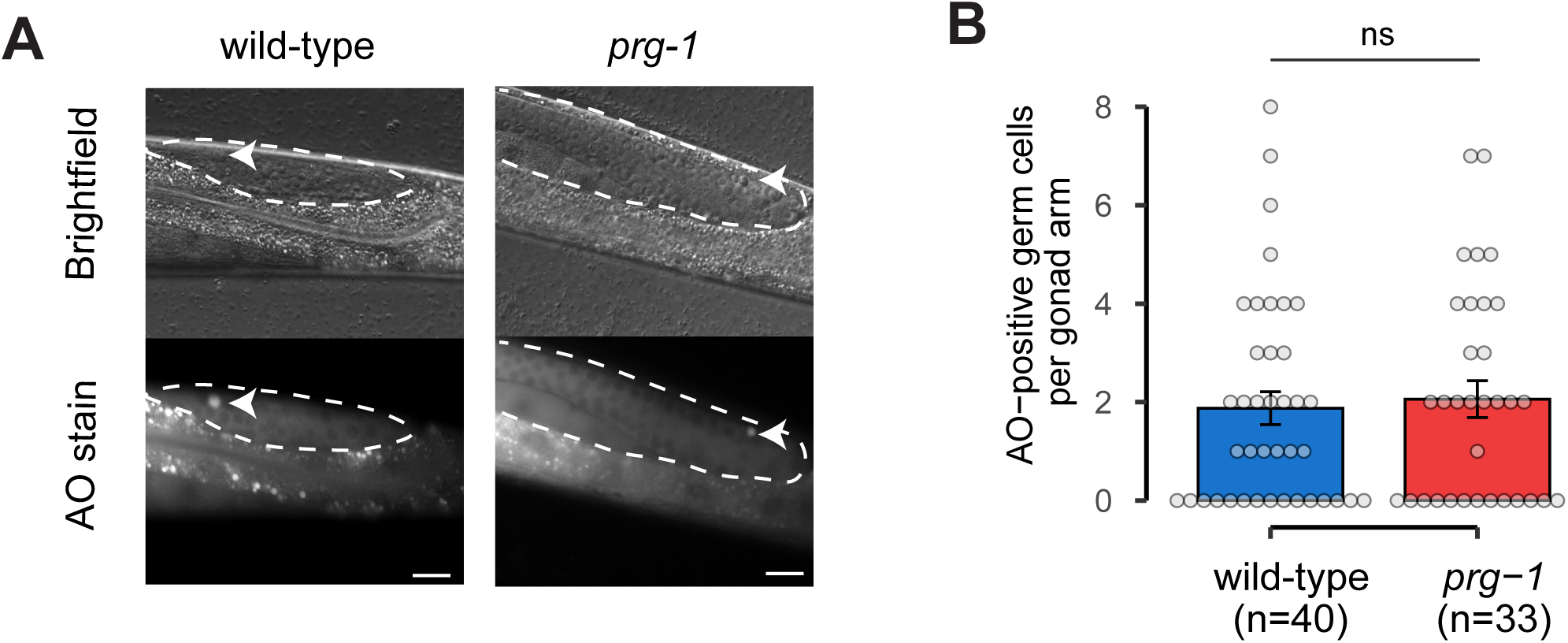
Acridine Orange staining of apoptotic corpses in the *C. briggsae* **germline.** (A) Representative brightfield and GFP fluorescence images of Acridine Orange (AO) stained germ lines of live adult wild-type and *prg-1* mutant animals under a 40x objective for the detection of germ cell apoptosis. Dashed lines outline the position of germline. Scale bar = 20 µm. (B) Bar plots quantifying the mean number of Acridine Orange (AO)-positive germ cells per gonad arm. in wild-type (blue, n=40) and *prg-1* mutant (red, n=33) animals. Each point represents a gonad arm assayed per animal. The Two-tailed Wilcoxon Rank Sum Test was used to compare number of AO corpses, ns represents p-value ≥ 0.05.

**Supplementary Figure 3:**
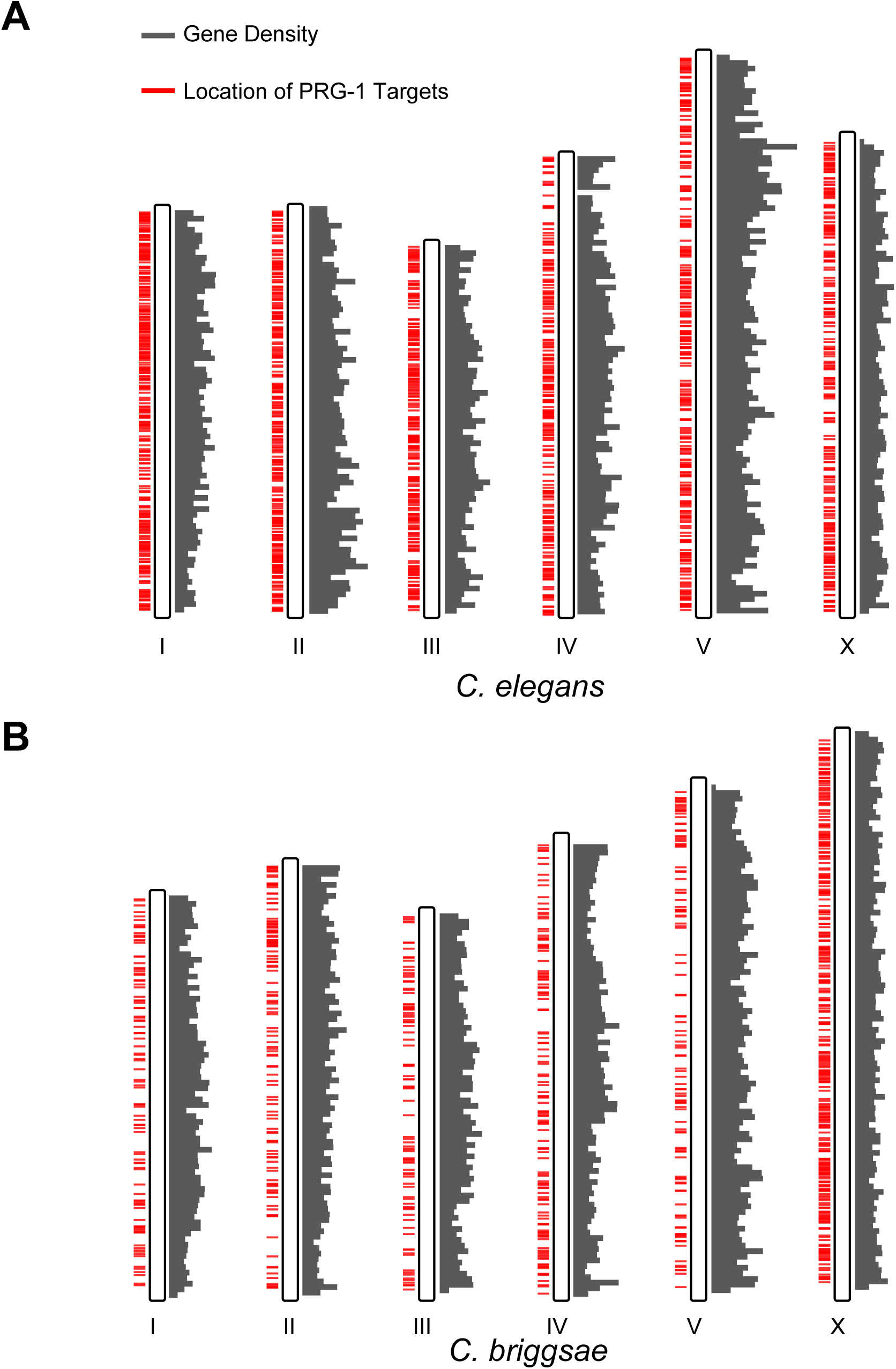
Genomic location of PRG-1 targets in *C. elegans* and *C. briggsae*. (A-B) Karyotype style plots showing the gene density (grey bars) as well as location of PRG-1 targets (red bars) along chromosomes I-X in *C. elegans* (A) and *C. briggsae* (B).

**Supplementary Figure 4:**
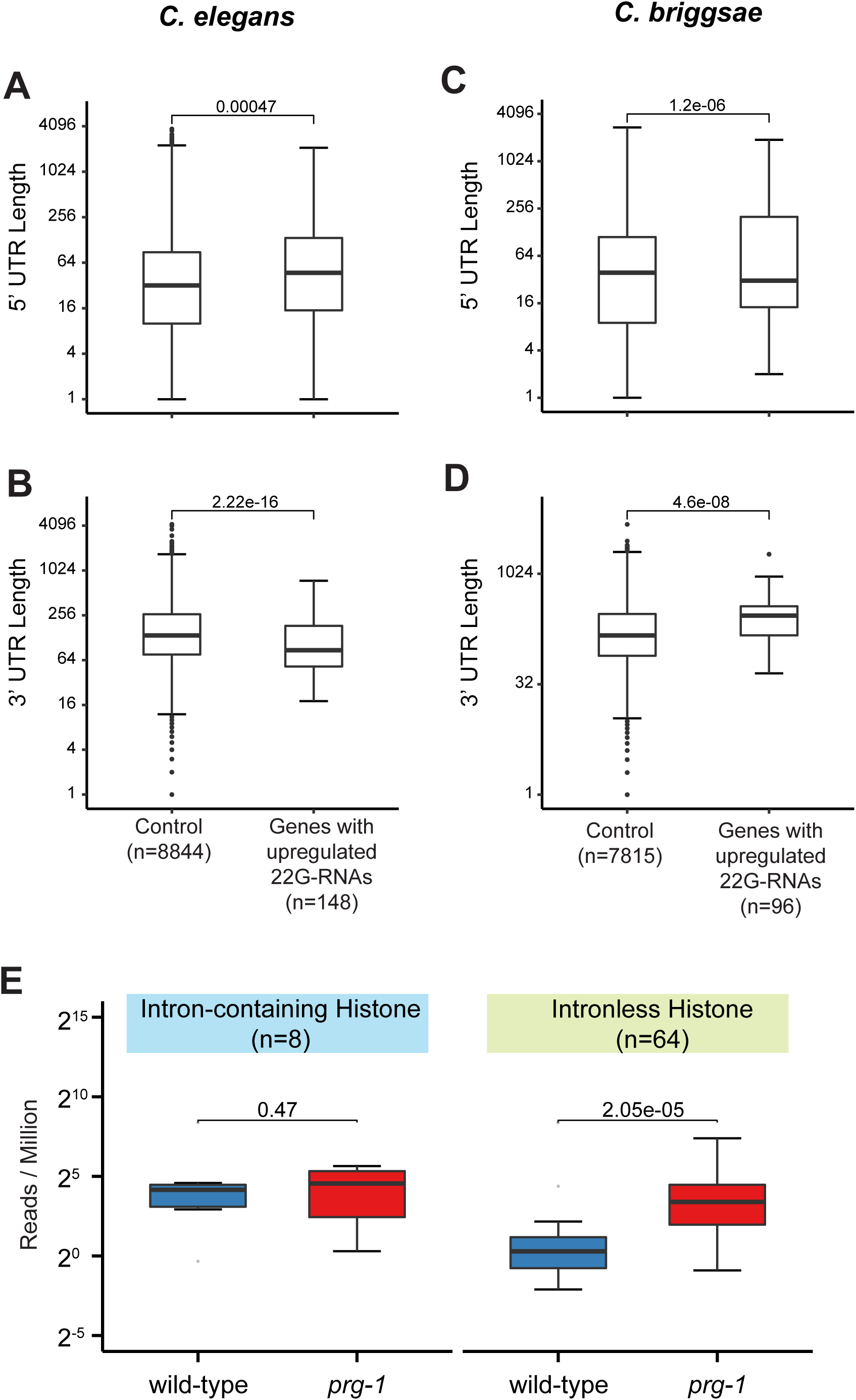
Length of 5’ and 3’ UTRs in *C. elegans* and *C. briggsae* for genes that accumulate 22G RNAs in *prg-1* mutants. (A-D) Boxplots showing the length of 5’ UTR (A and C) and 3’ UTR (B and D) in *C. elegans* (A and B) and *C. briggsae* (C and D) for transcripts that accumulate 22G-RNAs in *prg-1* mutants. The sample size of control and genes with upregulated 22G-RNAs in *C. elegans* and *C. briggsae* are denoted in x-axis labels. Genes in the control set are expressed in germline and do not exhibit changes in 22G-RNA levels in *prg-1* mutants. p-values were derived using a Two-tailed Wilcoxon Rank Sum Test. (E) Boxplot showing the abundance of 22G-RNAs targeting intronless or intron-containing histone genes in wild-type and *prg-1* mutant animals. P-values were derived using a Two-tailed t-test.

**Supplementary Table S1.** *C. briggsae* piRNA genes defined by the enrichment from Cbr-PRG-1 IP.

**Supplementary Table S2:** 22G-RNA density in *C. briggsae*.

**Supplementary Table S3:** 22G-RNA density in *C. elegans*.

**Supplementary Table S4:** Conserved PRG-1 Targets in *C. briggsae* and *C. elegans*.

**Supplementary Table S5:** Worm strains used in this study.

All Supplementary tables and metadata files are deposited to and are available at Dryad (https://datadryad.org/stash/share/JG5_VTsoNyi_62H03WHjnHH1wD2ZwZMLWpL4SnD1JTA)

